# Non-invasive laser speckle imaging of extra-embryonic blood vessels in intact few-days-old avian eggs

**DOI:** 10.1101/2024.03.11.584528

**Authors:** Zhenyu Dong, Simon Mahler, Carol Readhead, Xi Chen, Maya Dickson, Marianne E. Bronner, Changhuei Yang

## Abstract

Imaging blood vessels in early-stage avian embryos has a wide range of practical applications for developmental biology studies, drug and vaccine testing, and early sex determination. Optical imaging such as brightfield transmission imaging offers a compelling solution due to its safe non-ionizing radiation, and operational benefits. However, it comes with challenges such as eggshell opacity and light scattering. To address these, we have revisited an approach based on laser speckle contrast imaging (LSCI) and demonstrated a high quality, comprehensive and non-invasive visualization of blood vessels in few-days-old chicken eggs, with blood vessel as small as 100 µm in diameter (with LSCI profile full-width-at-half-maximum of 275 µm). We present its non-invasive use for monitoring blood flow, measuring the embryo’s heartbeat, and determining the embryo’s developmental stages using machine learning with 85% accuracy from stage HH15 to HH22. This method can potentially be used for non-invasive longitudinal studies of cardiovascular development and angiogenesis, as well as egg screening for the poultry industry.

## Introduction

Chicken and other avian embryos have served as fundamental research models in developmental biology as well as for the pharmaceutical industry, vaccine development, and agriculture. Of particular research interest is the chorioallantoic membrane (CAM), a vascularized extraembryonic membrane containing a network of blood vessels surrounding the chicken embryo that plays a crucial role in its development. Specifically, it facilitates the transportation of oxygen and nutrients, and assures protection of the developing embryo. Previous studies have shown that the CAM development can provide valuable insights on the cardiovascular system, and the vascularized embryonic membranes^1–4^. Additionally, CAM provides a unique biological microenvironment for evaluating tumor growth and angiogenesis^3,4^. Therefore, there is a continuous effort in developing methods for imaging the CAM.

Two primary methods have been widely employed to perform high resolution imaging of the CAM of avian embryos. The first is to culture the entire embryo outside the egg in vitro^5,6^. The second is to cut a hole in the eggshell, remove the protective membrane, and substitute it with a transparent window to access the embryo in ovo^7,8^. Common high resolution imaging techniques that can be used to image the exposed CAM include white light imaging^3^, fluorescence or confocal microscopy^4^ and optical coherence tomography^9,10^. As the process of exposing the CAM involves breaking the eggshell and potentially removing the outer and inner membranes of the embryo, the natural development of the embryo is altered. This alteration changes the gas exchange dynamics, cardiodynamics, physical tension on the embryo, and resistance to infection, negatively impacting the overall survival of the embryo^11,12^. Thus, having the ability to observe the CAM non-invasively within the intact egg, throughout early stages of embryonic development would be beneficial for various scientific studies, commercial applications, and conservation^11,13^. The challenge lies in developing an imaging method that enables visualization of the embryonic tissues such as the CAM, heart, and blood vessels while overcoming the opacity of the eggshell.

There are two non-invasive optical imaging methods that have been used to perform egg imaging. The first imaging method, brightfield transmission imaging, also commonly called egg candling, is the traditional optical method for observing the blood vessels within a chicken egg^17,20,21^. It is primarily used for assessing the viability of the embryo in agriculture and avian conservation^13,14^. A broadband light source is placed beneath or behind the egg. As the egg is held against the light source, the internal elements can be visualized due to the differences in the absorption of the light^13–15^. As egg candling relies on absorption of the illuminating light by the sample, the imaging can be compromised by undesired external sources of light absorption^15^, such as cracks or pigmentation of the eggshell as well as air-bubbles, especially between the inner and outer shell membranes. In addition, precise control of the illumination power is imperative to prevent issues of overexposure and underexposure^15^.

The second non-invasive imaging method, laser speckle contrast imaging (LSCI), relies on the interaction between scattered laser light and dynamic samples, such as blood cells^16–19^. These interactions create speckle patterns, which can be recorded and used for reconstructing the image of the sample. LSCI has gained significant interest for visualizing blood vessels in a diverse types of living tissues^16–22^, including the chicken egg^23–26^. For example, LSCI has been applied for imaging chicken embryos through intact egg at early incubation stages^26^. Results published in the paper^26^ showcased imaging of the vascular structure of the chicken embryo during the first few days (day 2 to day 5) of incubation. However, starting from the fourth day of incubation, the quality of the image became obscured, due to the increased presence of surrounding scattering tissues. Indeed, LSCI imaging is contingent on the dynamics of the sample and is compromised by the presence of undesirable dynamic elements in the surroundings. Furthermore, the temporal resolution reported in the paper was insufficient to monitor blood flow dynamics or heartbeat of the embryo. Lastly, to achieve optimal imaging performance, precise prior alignment of the sample was required to ensure the accurate focus on blood vessels.

To address these challenges, we have implemented an improved LSCI system that is capable of imaging the blood vessel network of few-days-old chicken eggs using temporal Fourier filtering and additional speckle noise reduction techniques^27,28^. Our results showed a significant improvement for imaging the chicken egg blood vessel network than previously reported^26^. For this study, we implemented both an LSCI system and a brightfield transmission imaging system on a single platform so that we can compare the performance of both methods. This paper is structured as follows: first, we present and discuss the strengths and limitations of each imaging method individually. Second, we report a comprehensive, non-invasive visualization of blood vessels of avians eggs with good spatial and temporal resolution. Third, we demonstrate some practical applications of our LSCI imaging system. Specifically, we have shown that our technique can be used to perform longitudinal tracking of blood vessel network development. In addition, we non-invasively monitored and visualized blood flow dynamics, and estimated the corresponding heartbeat within the chicken egg. Finally, we performed physical feature extraction on the blood vessel images obtained from LSCI and embedded them into machine learning algorithms to discern the embryonic development stages of the chicken embryo across four different Hamilton-Hamburger (HH) stage periods^29^. Our results provide a developmental stage classification accuracy of 85% based on a dataset consisting of 260 blood vessel images.

## Results

### System design and principles

To compare the light-emitting diode (LED)-based brightfield transmission imaging and laser speckle contrast imaging (LSCI), we combined the two imaging methods in a dual imaging platform (Fig. 1). Figure 1a shows the components of brightfield transmission imaging, comprised of an LED for illumination and a camera for detection. In our configuration, a 530 nm LED illuminated the egg from underneath; and the incident light was absorbed by various components (yolk, embryo, CAM, eggshell) and the transmission image was collected by an RGB camera. The absorption results in emission of a fluorescence signal of longer wavelength (see Supplementary Fig. 2), typically above 600 nm^30,31^. Consequently, as shown in Fig. 1a, the red channel of the camera captured the fluorescence image, while the green channel captured the absorption image. As anticipated, the blue channel of the camera did not detect any appreciable signal.

**Fig. 1.**
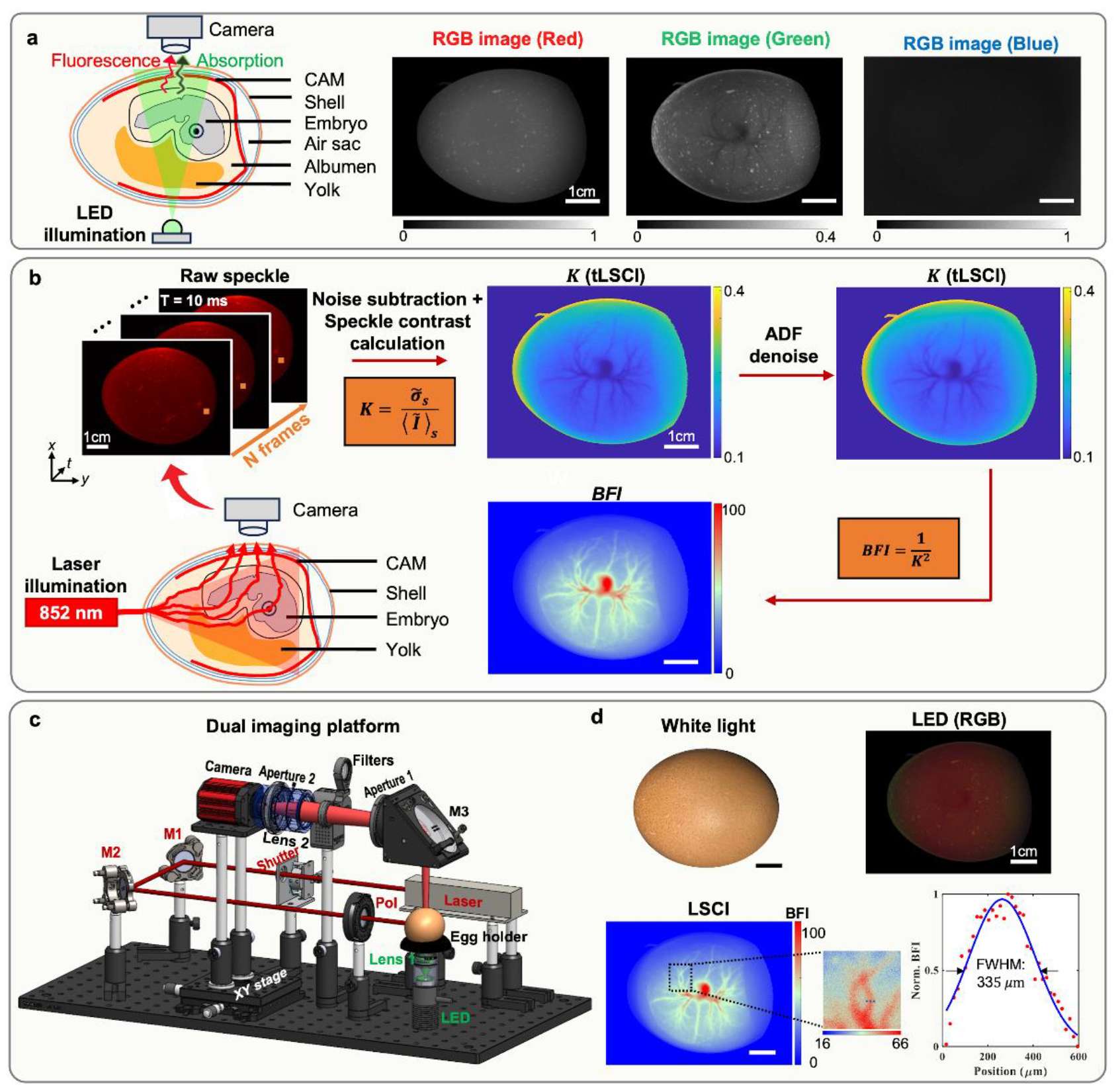
Experimental arrangement of the brightfield transmission imaging and the laser speckle contrast imaging for visualizing the blood vessels of a chicken egg. **a** Brightfield transmission imaging. The green channel image was contrast enhanced for visualization. **b** LSCI imaging method and reconstruction pipeline. **c** Dual imaging platform combining LED brightfield transmission imaging and LSCI imaging. M1, M2, M3: mirrors. Pol: linear polarizer. **d** Typical imaging results. The normalized BFI profile along the blue dotted line in the zoomed LSCI image represents a full width at half maximum (FWHM) of 335 µm.

LSCI is a well-established method for visualizing blood vessels in diverse types of living tissues^16–21^, including chicken eggs^23–26^. Its non-invasive nature and capacity to capture dynamic physiological processes make LSCI a valuable tool in the study of vascular dynamics and blood flow patterns, contributing to a deeper understanding of circulatory systems in various biological contexts. When a coherent laser beam is directed onto the sample (i.e., a chicken egg), light will experience multiple random scattering events before exiting the sample. In our case, we illuminated the chicken egg from the side and collected the exiting light from the top of the egg, see Fig. 1b. As opposed to brightfield transmission imaging, the collected light was granular in appearance, also called speckles. Speckles arise from the mutual interference of light following different trajectories, caused by the random scattering of light in the sample^32^. As components (such as blood cells) within the sample move, the speckle field undergoes dynamical changes^33,34^. These speckle fluctuations can be characterized by the decorrelation time *τ*_*c*_ of the speckle field^17,35^. This decorrelation time is related to the inherent dynamics of the sample. The dynamic components in the egg and their rate of change can be visualized and quantified by calculating the speckle contrast value *K* (see Methods) of speckle patterns recorded by the camera^35^, defined as the ratio between the standard deviation and the mean value of the pixels intensities. To achieve this, the camera exposure time *T* should be significantly longer than the decorrelation time *τ*_*c*_, ensuring that the recorded speckle patterns represent an ensemble average of the decorrelation events. As the moving blood cells in the vessels exhibit greater movements than the yolk and other background tissues (i.e. shorter decorrelation time), the speckle contrast of the blood vessels can be expected to be lower than the rest of the area.

In Fig. 1b, we outline our processing pipeline for LSCI imaging. The camera recorded a sequence of *N* = 200 speckle frames images when the laser was on and another *N* = 200 noise frames when the laser was off with the same exposure time, see Methods for additional details. The recorded noise frames were then subtracted from the speckle frames. The speckle contrast values were calculated for each camera pixel in the temporal domain over the *N* noise subtracted speckle frames, which is commonly denoted temporal LSCI (tLSCI)^20,22^. As blood cells move within the blood vessels, this movement generated multiple speckle realizations within the camera exposure time, leading to a smeared and washed-out speckle pattern recorded by the camera. The smearing effect, due to the dynamics of the blood cells, leads to lower temporal fluctuations, decreasing the speckle contrast value. In this study, we chose to use tLSCI rather than spatial LSCI (sLSCI) which employs a sliding window in the spatial domain^17^, since tLSCI can help to remove the static scattering such as cracks on the eggshell, see Supplementary Fig. 3 for more details. To reduce the speckle noise after tLSCI reconstruction, we used an anisotropic diffusion filter (ADF), which can preserve image details while enabling image smoothing^27,28^. We also found that the application of ADF denoising enhanced the temporal resolution of the imaging, achieving imaging of the blood vessels with as few as *N* = 3 speckle frames, see additional results in Supplementary Fig. 3. Finally, as shown in Fig. 1b, we converted the reciprocal of squared speckle contrast *K*^2^ to blood flow index (BFI) to quantify the blood flow, which is inversely related to the decorrelation time *τ*_*c*_ of the blood flow dynamics (see Methods)^36^. More details regarding the LSCI equations can be found in the Methods section.

In Fig. 1c, we show a dual imaging platform, built to compare the two imaging methods. The brightfield transmission imaging platform (green legend) was comprised of a green LED of central wavelength of 530 nm (480 mW) illuminating the bottom of the egg. The LSCI imaging platform (red legend in Fig. 1c) used an 852 nm near-infrared laser (230 mW) to produce side illumination onto the egg, allowing high photon transmission efficiency through the egg. The two methods shared the same detection configuration (i.e., the same camera), where the top-view of the egg was imaged by an RGB camera with a tunable lens for focus adjustment. An adjustable aperture was placed in the pupil plane of the lens to enable adjustment of the speckle size of the light field in LSCI. It was important to adjust the speckle size to be close to two pixels per speckle, for precise speckle contrast estimation^37–39^. The magnification of the imaging system was 0.2 and the equivalent numerical aperture was 0.03. Detailed descriptions of the experimental setup can be found in the Methods section.

Figure 1d shows typical imaging results from the brightfield transmission imaging and LSCI imaging. Notably, LSCI imaging is of superior quality and resolution, providing clear visualization of the blood vessel network, encompassing vessels of varying size. As shown in the zoomed image, smaller blood vessels can also be visualized. The cross-section of the selected blood vessel in the zoomed image (blue dashed line) shows the capability of our method to measure the profile of a blood vessel.

In order to determine the smallest blood vessel diameter detectable by our LSCI platform, we compared LSCI images with microscope images obtained after opening the egg and removing the protective membrane, see Fig. 2. Across five different eggs, we selected the three smallest blood vessels visible in the LSCI image and measured their average full width at half maximum (FWHM) across the five eggs to be 275 ± 25 µm. In the corresponding microscope images, we measured the average diameter of the selected blood vessels across the five eggs to be 100 ± 12 µm. This set out the smallest blood vessel diameter detectable by our LSCI platform to be 100 µm. The difference between the two measurements is attributable to the point spread function of the transmitted scattered light field as collected by the LSCI imaging system. The light scattered from the blood vessels is further scattered by the eggshell upon exiting the egg. Both of these processes contribute to the degradation of the attainable resolution.

**Fig. 2.**
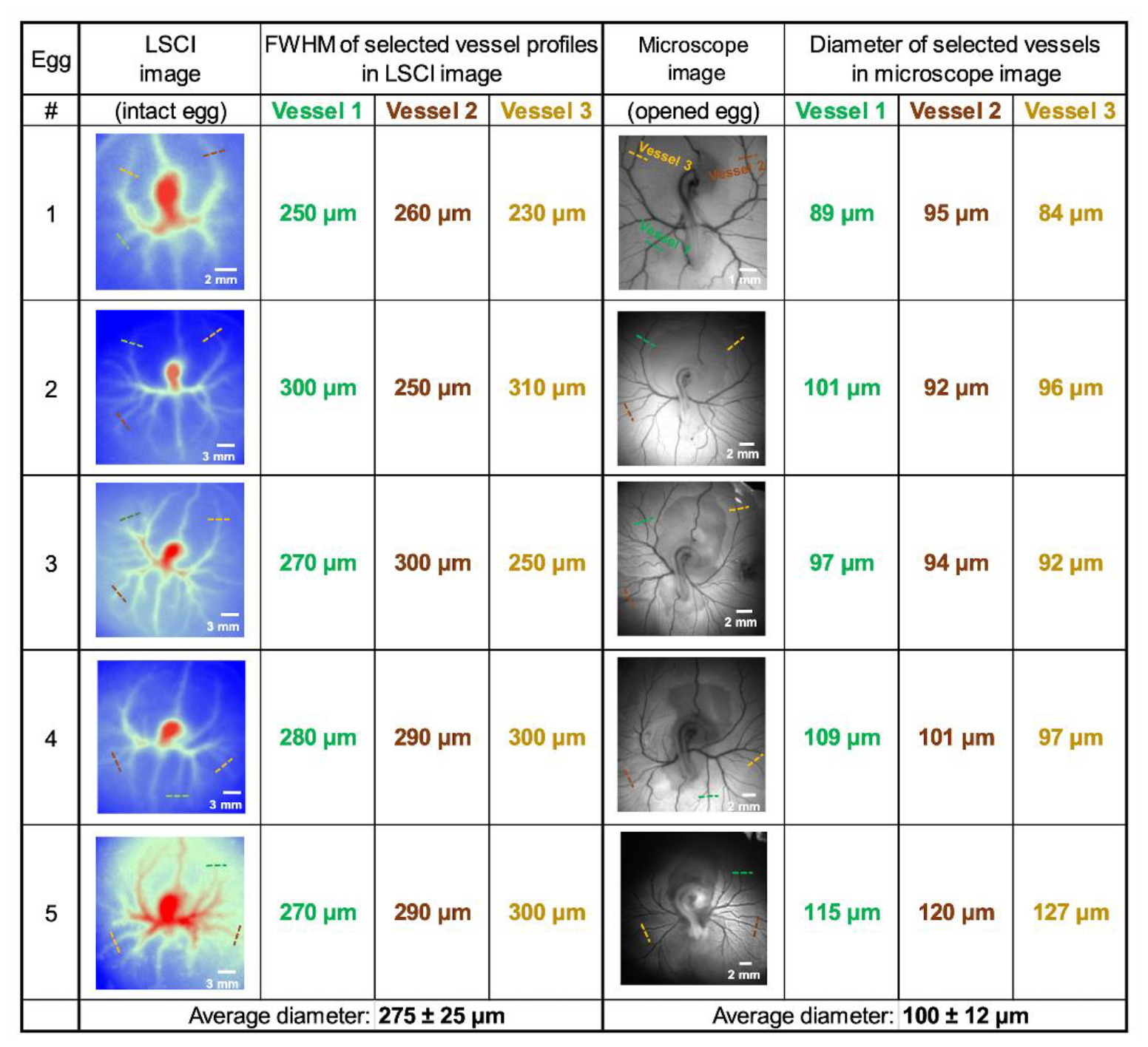
Smallest blood vessel diameter detectable by the LSCI platform in chicken eggs. Left panel: LSCI images of five different eggs. FWHM: full width at half maximum. The three smallest detectable blood vessels in the LSCI image were selected and the FWHM of their profiles were measured. Right panel: Microscope images after opening these eggs. The selected three smallest visible blood vessels were identified, and their diameters were measured from the microscope image.

### Comparison between brightfield transmission imaging and LSCI

To compare the brightfield transmission imaging and LSCI methods, we imaged six chicken eggs at day 4 (96 hours) of incubation. See Methods for details about our egg incubation procedure. Figure 3a shows photographic images of each egg under white light illumination. The six chicken eggs were selected as follows: one brown infertile egg, one white fertile egg, and four brown fertile eggs with varying degrees of brown coloration, labeled as Brown 1 through Brown 4, according to their shade (from bright to dark).

Figure 3b shows the brightfield transmission imaging results. The first egg (labeled ‘Infertile’) did not contain any blood vessels, indicating a lack of embryo development. Notably, the yolk within the egg was visible, confirming its infertility. For the white egg, the blood vessels surrounding the embryo were distinctly visible. However, for the brown eggs, the visibility of the blood vessels diminished as the eggshell had darker colorations. The blood vessels were readily observable in Brown 1 and Brown 2, but became less discernible in Brown 3, and were not visible in Brown 4. These findings showed that the color of the eggshell has a significant impact on the brightfield transmission imaging results. The imaging was primarily influenced by the absorption phenomenon, as depicted in Fig. 1a. When the eggshell exhibits a darker coloration, the absorption effect becomes increasingly dominant, adversely affecting the imaging of blood vessels (see the absorption line profiles in Fig. 3b).

Figure 3c shows the LSCI imaging results. For the infertile egg, only the movement of the yolk was observed, characterized by a relatively low BFI. In contrast, both white and brown eggs exhibited distinct visibility of blood vessels surrounding the embryo and shared similar BFI values. Notably, there was no disparity in LSCI imaging quality across white egg, dark brown egg (e.g., Brown 3), and light brown egg (e.g., Brown 1), as shown in the BFI line profiles in Fig. 3c, where the position of the blood vessels can be identified by the peaks consistently. These observations underscore that the color of the eggshell has no significant impact on the LSCI imaging results. This phenomenon can be attributed to the operating principle of LSCI imaging, which hinges primarily on speckle dynamics, specifically the motion within the imaged sample. Since the eggshell remains largely static (non-moving), the speckle pattern originating from the eggshell is stationary (static speckle pattern) and is effectively filtered out during temporal speckle contrast calculations. Conversely, as there are blood flow movements in the vessels of the chicken embryo, the speckle pattern undergoes changes over time (resulting in a dynamic, moving speckle pattern) and is captured by higher BFI values through temporal LSCI computations. Consequently, eggshell cracks were apparent with the brightfield transmission imaging method (see dip positions in the absorption line profiles in Fig. 3b) but were absent with the LSCI method (Fig. 3c). This highlights a significant advantage of the LSCI method over brightfield transmission imaging.

To confirm our findings, we repeated the experiment of Fig. 3 (at day 4 of incubation) using a set of six quail eggs, comprising one infertile quail egg and five fertile quail eggs, see Fig. 4. The quail eggs, smaller than the chicken eggs, each featured a white eggshell adorned with random black patches. The results obtained from the quail eggs are particularly noteworthy. The blood vessels can be readily observed on white regions of the eggshell, whereas they are not visible on eggshell’s regions with black spots. The LSCI imaging performs well on all eggs, regardless of the eggshell’s coloration.

Based on the results reported in this section, we can see that the LSCI method consistently overperforms the brightfield transmission imaging method. In the following three sections, we report on our study of specific LSCI imaging performances and demonstrate a number of practical applications of our LSCI imaging system.

### Longitudinal imaging of embryonic blood vessel network development

In this section, we demonstrate the ability of the LSCI imaging system to track blood vessel network growth. To this end, we partitioned our imaging sessions based on the incubation time. As shown in Fig. 5, we imaged both a white and a brown chicken egg across three incubation days: day 3 (72 hours), day 4 (96 hours), and day 5 (120 hours). Between day 3 and day 5, both the white (Fig. 5a) and brown (Fig. 5b) eggs exhibit a progressive expansion of the vessel network and an increase in the BFI.

On day 5 (120 hours) of incubation, the blood vessels of the chicken embryo undergo a significant increase in density and begin to develop more deeply within the egg, leading to more absorption of light and decreased number of detectable photons on the camera. Under this condition, the LSCI method is more sensitive to light scattering from the shell and to the illumination intensity fluctuations caused by embryo body movements (see Supplementary Fig. 4 for more details). Therefore, in LSCI images, spot-shaped imaging artifacts become evident, matching the crack position of the shell (black arrows in third column of Fig. 5a and Fig. 5b). In addition, as the embryo advances to a later stage of development, increased movement of the embryo within the egg becomes notable. Thus, at day 5, movement artifacts during our acquisition time create a non-uniform background and reduced contrast of the blood vessels in LSCI imaging.

These two imaging artifacts resulted in low-frequency signals that can be filtered out by devising a time domain Fourier high-pass filter, as depicted in Fig. 5. Further details of the time domain Fourier high-pass filtering can be found in Supplementary Fig. 4. As evident, the filter effectively suppresses the structured dot artifacts and the uneven noise background, and the contrast of the blood vessels with respect to the background is significantly enhanced. We were ultimately able to obtain a clear blood vessel image with higher contrast and fewer artifacts compared to before. As shown by the line profiles, a larger peak-to-valley range is observable in which sub-branches of blood vessels can be resolved (red arrows).

By using Fourier filtering, a significant improvement has been achieved for imaging the chicken embryo blood vessel network than previously reported^26^.

### Non-invasive monitoring of the blood flow dynamics with a chicken embryo at early stage of incubation

The results for monitoring the blood flow in a chicken egg non-invasively at day 4.75 (114 hours) of incubation are presented in Fig. 6. To monitor the blood flow variations as a function of time, we employed the LSCI imaging method with *N* = 300 recorded speckle pattern images. Instead of performing LSCI with a temporal speckle contrast calculation across all *N* = 300 images, as in equation (7) in Methods section, we used a sliding window in time domain over *n* = 3 adjacent speckle pattern images to create a series of blood flow reconstructions over time, see Supplementary Fig. 1b. It is also shown in Supplementary Fig. 3 that with ADF denoising, three adjacent frames can provide a reasonable estimate of the blood flow.

The RGB camera operated at 21 frames per second (FPS), resulting in a movie of the chicken embryo’s blood flow distribution map at 21 images per second (see Fig. 6a and Supplementary Movie). The rectangular sliding window in time domain acts as a 7 Hz low-pass filter in the frequency domain, whose temporal Fourier transform is a Sinc function. Previous studies have shown that chicken embryos between day 3 and day 5 of development typically have a characteristic heartbeat frequency between 2 and 3 Hz^40^. According to the Nyquist sampling theorem, our method can acquire information associated with the first and second harmonics of the heartbeat event. Using the sliding window method, we can obtain real time imaging of the blood vessels, as shown in the Supplementary Movie. Such live view ability helped us to swiftly align the sample to its best possible imaging position (i.e. to place the embryo in the center of the image) before starting the recording, ensuring optimal performance.

Figure 6a shows three snapshots from the LSCI movie. Each snapshot corresponds to a specific time of the cardiac cycle of the chicken egg, see Fig. 6b colored boxes for time indication. See Supplementary Movie for the full video. As expected, the blood flow at different periods of a cardiac cycle can be observed. During systole, the heart contracts, propelling blood into the arteries, leading to high BFI. In contrast, during diastole, the heart relaxes and refills with blood, causing a reduction in BFI within the arteries.

Figure 6b shows the averaged BFI value within the center cropped region calculated from each image of the movie shown in Fig. 6a. The heart rate was calculated by Fourier transforming the blood flow signal, see right panel of Fig. 6b. A hamming window was applied before the Fourier transform to avoid spectral leakage. The Fourier amplitude peak centered around *freq*_*HR*_ = 2.43 Hz corresponds to the heart rate amplitude peak of the embryo. There was also a Fourier peak centered at 4.9 Hz corresponding to the second harmonic of the heart rate pulsations of the embryo^41^.

Figure 6c shows the temporal dynamics of the blood flow measured by using the speckle visibility spectroscopy (SVS) method^41,42^. In SVS, instead of calculating the blood flow from the temporal speckle contrast of three adjacent speckle pattern images, the blood flow value at one time point was calculated from speckle contrast of each speckle pattern image, with each camera pixel acting as a sample point^42^ (see Supplementary Fig. 1c). As shown, similar results were obtained as with the LSCI calculation method. However, the SVS results were slightly noisier than the results in Fig. 6a, since SVS typically require a higher sampling rate (above 30 FPS) and cannot eliminate the influence of static scattering without pre-calibration of the noise^41–43^. We anticipate that SVS results would show better signals within a cardiac cycle if one were to use a camera with an FPS of 50 or more. The Fourier transform of the SVS signal shows Fourier peaks at the same frequencies as in Fig. 6a, confirming the accuracy of our heartbeat frequency estimation using LSCI. However, SVS cannot provide high-resolution spatial distribution of blood flow, which is the reason why we prefer using the LSCI method here.

Figure 6d shows the temporal dynamics of the blood flow measured after opening the egg, by recording microscope camera images of the embryo (see Supplementary Movie), using the technique of Photoplethysmography (PPG)^44^. The intensity fluctuations here reflect absorption changes due to the embryo cardiac circulation. Blood flow measurements closely resemble those obtained using the LSCI or SVS methods, but with an enhanced signal quality, attributable to the direct access to the embryo achieved by opening the egg. The Fourier transform reveals two distinct peaks. The first of *freq*_*HR*_ = 2.62 Hz, corresponds to the amplitude peak of the embryo’s heart rate and the second peak at 5.24 Hz to the second harmonic. This result confirms that the LSCI method can be used to measure the heartbeat non-invasively with good fidelity.

### Staging chicken embryo development via blood vessel image classification

In this section, we aim to demonstrate that the LSCI images when used with machine learning algorithm can be effective in the classification of the developmental stages of the early chicken embryo. Accurate staging of chicken embryos is important for early developmental biology studies, particularly in monitoring the development of organs such as the heart, limbs, retina, and other vital structures^45,46^. Performing this task is challenging, typically involving the delicate process of opening the eggshell and membrane, followed by a meticulous examination of the embryo conducted by experienced biologists^29^. By performing staging non-invasively with LSCI, we can not only avoid the manually intensive task of eggshell removal but also enable the possibility of longitudinal studies with the same egg.

Figure 7a displays a segment of the widely recognized Hamburger-Hamilton (HH) staging series^29^ captured with a wide-field microscope. This series has established itself as a gold standard in biology literature for precise staging results as it offers a direct visual representation of the chicken embryo’s morphological growth. Essential features such as the tail’s shape, the head’s direction, lateral body-folds, and the limb development, collectively contribute to discerning differences between each stage^29^. Here, we want to show that with our non-invasive imaging technique (LSCI), we can perform quantitative analyses on the blood vessel distributions and distinguish between stages, a departure from the traditional HH staging method relying on morphological differences observed from the embryo body after opening the egg.

In our experimental design, we focused on four paired stage periods: HH15-16, HH17-18, HH19-20, and HH21-22, where our LSCI system produces high-quality images. The development of a chicken embryo is a continuous process, which is commonly studied by grouping adjacent HH-stages together (stage periods), such as in heart studies^47^. The period from stage HH15 to HH22 marks a fast-growing period in blood vessel expansion, consequently, achieving accurate staging is particularly useful, enabling in-depth studies of each stage or facilitating data normalization within the same developmental range. In Fig. 7b, we compared the non-invasive blood flow map using the LSCI method with the microscope image after opening the egg. The microscope image reveals the complete structure of the CAM and the shape of embryo body. To quantify the vessel characteristics, we performed feature extraction using image processing methods and referred to the result as LSCI+. This method can automatically segment out blood vessel regions together with BFI value, vessel skeleton and branch points as quantitative outputs. The LSCI+ and microscope images in Fig. 7b show highly relevant details of the blood vessel distribution among all the four stages. More details about the microscope images acquisition and the image processing pipeline can be found in Methods and Supplementary Fig. 5.

To train the machine learning algorithms to perform classification, we collected a dataset of 260 brown eggs in total, with 65 eggs for each stage. From our LSCI+ images, we extracted four features including vessel length, number of branches, vessel area, and standard deviation of BFI (Fig. 7c). We chose to use the normalized standard deviation of BFI value instead of the direct blood flow reconstruction value since the latter depends not only on developmental stage but also on the temperature and humidity which can be variable^26^. The data distributions of the four extracted features are shown in Supplementary Fig. 6, which indicates that using the feature space constructed by these four variables enables separation of the four stages. Since these features have physical meanings, this helps reduce the overfitting problem that convolutional neural networks usually suffer from when dealing with a small dataset.

The dataset was first randomly split into a training set (160 eggs) and a testing set (100 eggs), which were equally balanced among the four stages. The ground-truth stage of each egg was determined by using the traditional HH staging method on recorded microscope images of the chicken embryo, captured after opening the egg, see Methods for additional details. To ensure reliable classification results with this small dataset, we performed five-fold cross-validation on our training set to select the best machine learning model and parameters for our task (Fig. 7d). We tried seven traditional machine learning algorithms which are designed for multi-class classification tasks, including Decision Tree, Light Gradient-Boosting Machine (LGBM), K-Nearest Neighbors (KNN), Random Forest, Logistic Regression, Naïve Bayes, and Support Vector Classifier (SVC). We chose the SVC method as our target classifier since it outperformed the other methods and has the highest average accuracy of 81.3% on the validation set. We finally trained the SVC model again using the whole training set and performed testing of our method using the testing set which has remained quarantined and unseen by the training process. The results were evaluated using confusion matrix and average testing accuracy among four stages. The confusion matrix in Fig. 7e shows that the prediction of the algorithm on stages HH15-16 has the highest accuracy, while the predictions on the three other stages have misclassifications in the adjacent stages. Figure 7f shows that we achieved an average accuracy of 85% using the SVC algorithm on LSCI images. Additionally, we also conducted three human tests, which produced an average accuracy of 77%, see Supplementary Fig. 7 for confusion matrix comparison. Our results indicate that our method can be applied for automated and non-invasive staging of chicken embryos for stages HH15 to HH22.

Figure 7g presents a 2D visualization of the data point distributions in feature space consisting of the first two principal components using principal component analysis. The four stages have a rough boundary among their clustering and most of the wrongly classified cases result from the edge points or exceptional interior points. This indicates that the classification error may contribute to mislabeling by biologists because of indistinguishable microscope images, overlapping distribution of features, small dataset size, imprecise feature extraction, as well as lack of nonlinearity of the classification algorithm.

## Discussion

In summary, we have demonstrated that our LSCI imaging system is capable of non-invasively imaging the heart and the blood vessel network in avian embryos (such as chicken or quail), specifically focusing on early-stage chicken embryos. Our LSCI prototype is able to detect blood vessels of diameter as small as 100 µm. The scattering of light within the egg and the eggshell does have a resolution degrading effect on the image. We empirically observed that blood vessels of diameter of 100 µm are rendered with profile FWHM of 275 µm in the LSCI images (Fig. 2). Our observations highlighted the advantages of LSCI over brightfield transmission imaging, where LSCI does not only provide specificity in capturing blood flow information but is also resilient against variations in eggshell color and shade (Fig. 3, 4). Using our LSCI system, we conducted longitudinal imaging from day 3 to day 5 of incubation (Fig. 5a, b), which followed the development of the embryonic blood vessel network. By incorporating an ADF denoiser and the Fourier filtering method, we have enhanced the image quality of blood vessels within the chicken egg (Fig. 5a, b, Supplementary Fig. 4), surpassing the image contrast observed in previous studies using LSCI.

**Fig. 3.**
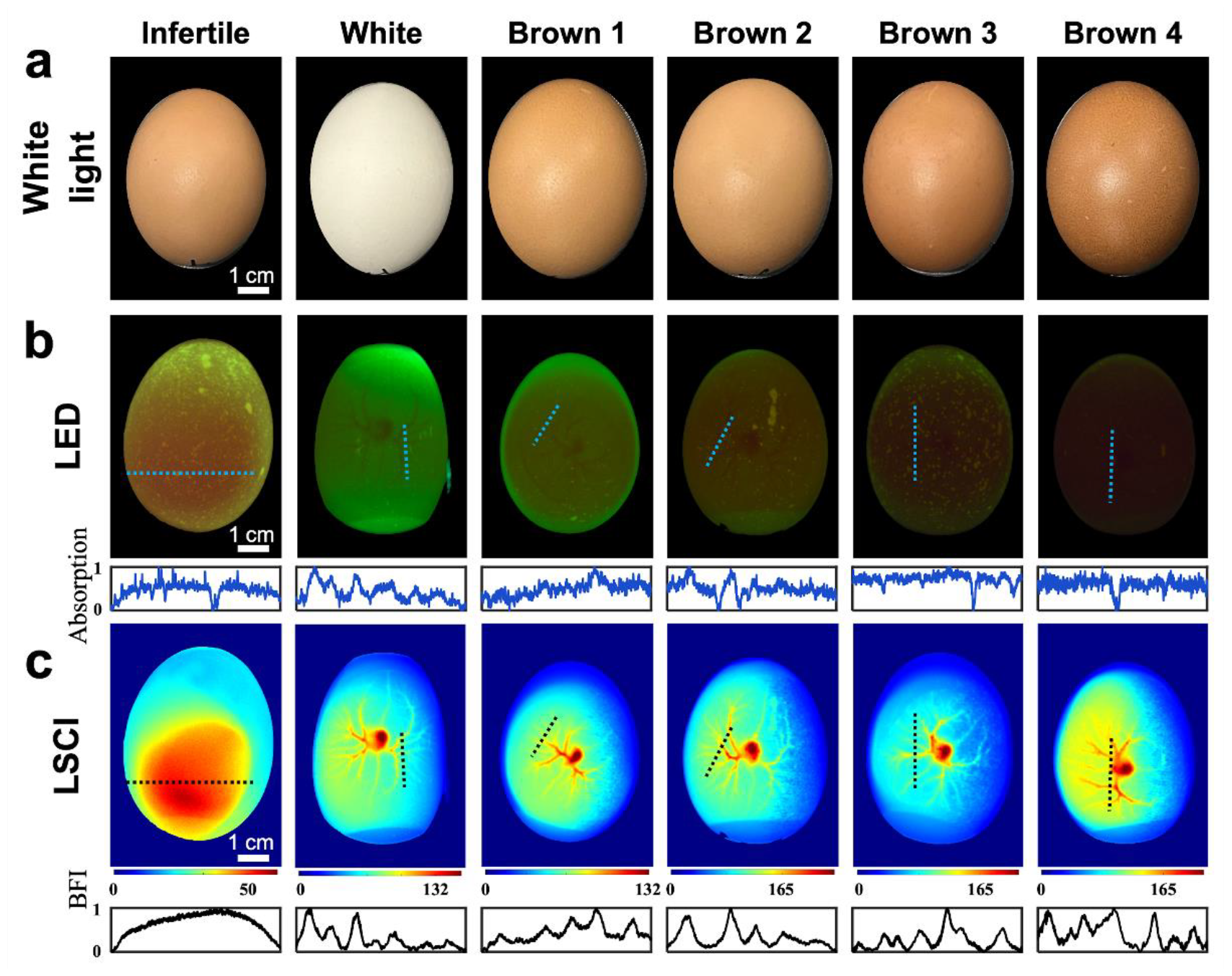
Blood vessel imaging in chicken eggs at day 4 of incubation. **a** Photographic image of each egg showcasing the coloration of the shell. **b** Brightfield transmission imaging method. The blood vessels in the white egg or light brown eggs (Brown 1 and Brown 2) can be observed but they cannot be observed in dark brown eggs (Brown 3 and Brown 4). The absorption line profiles are calculated as the inversion of normalized green channel intensity of the RGB images. The eggshell crack positions have lower absorption and show up as dips in the line profiles. **c** LSCI imaging method. The blood vessels can be effectively observed for all egg types. Note that the first egg is infertile, displaying no signs of an embryo. The BFI line profiles are normalized to between 0 and 1.

**Fig. 4.**
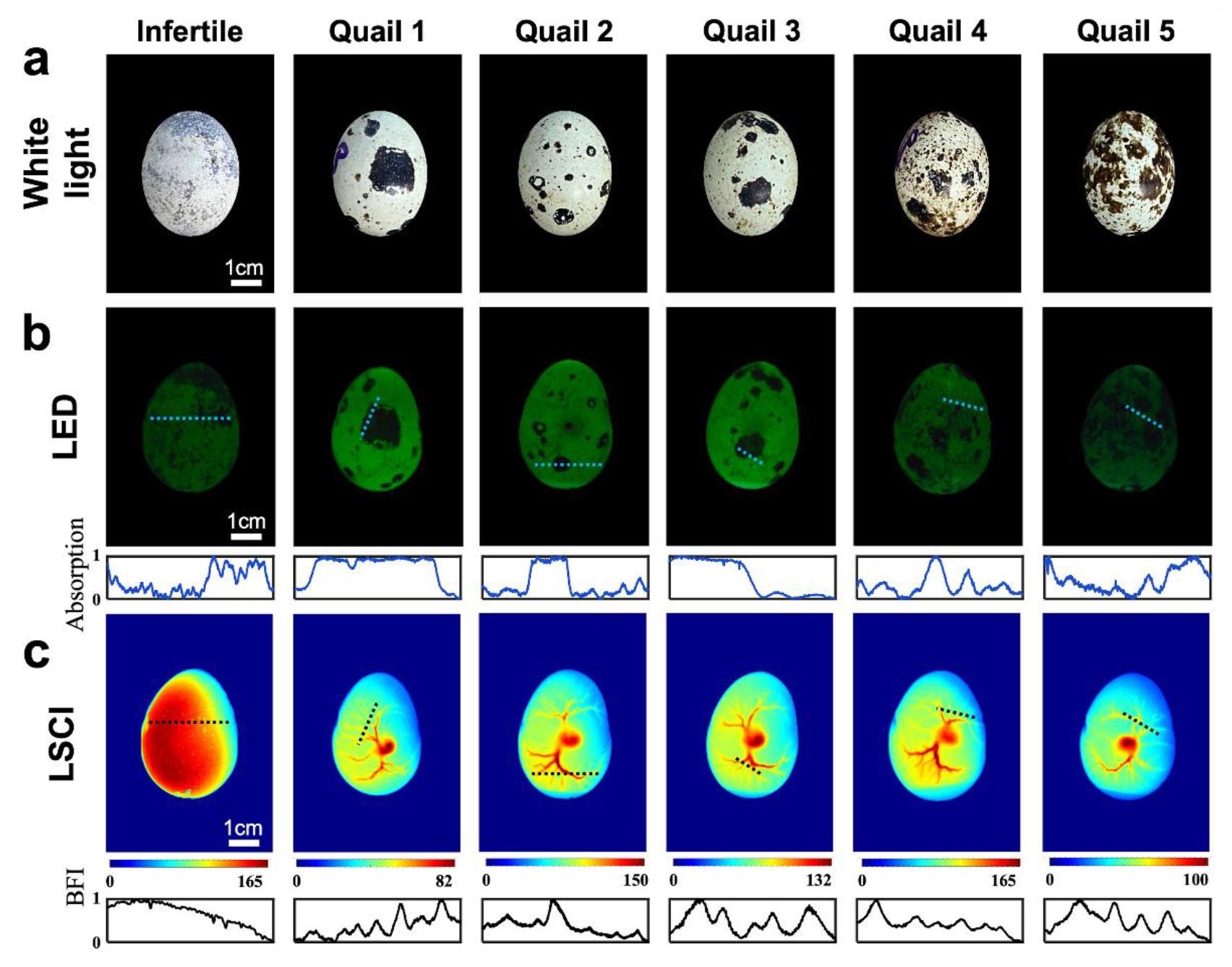
Quail eggs imaging with our dual imaging platform (day 4 of incubation). **(a)** Photographic image of each quail egg showing its coloration. **(b)** Brightfield transmission imaging results. Blood vessels are readily observable on white regions of the eggshell, but not on the black stained regions. The absorption line profiles show the high absorption in black stained regions of the eggshell. **(c)** LSCI imaging results. The BFI line profiles are normalized to between 0 and 1. Opposed to the brightfield transmission imaging method, the LSCI method is not influenced by the color, shade, or cracks of the eggshell. Note that the first quail egg is infertile, displaying no signs of embryo development.

**Fig. 5.**
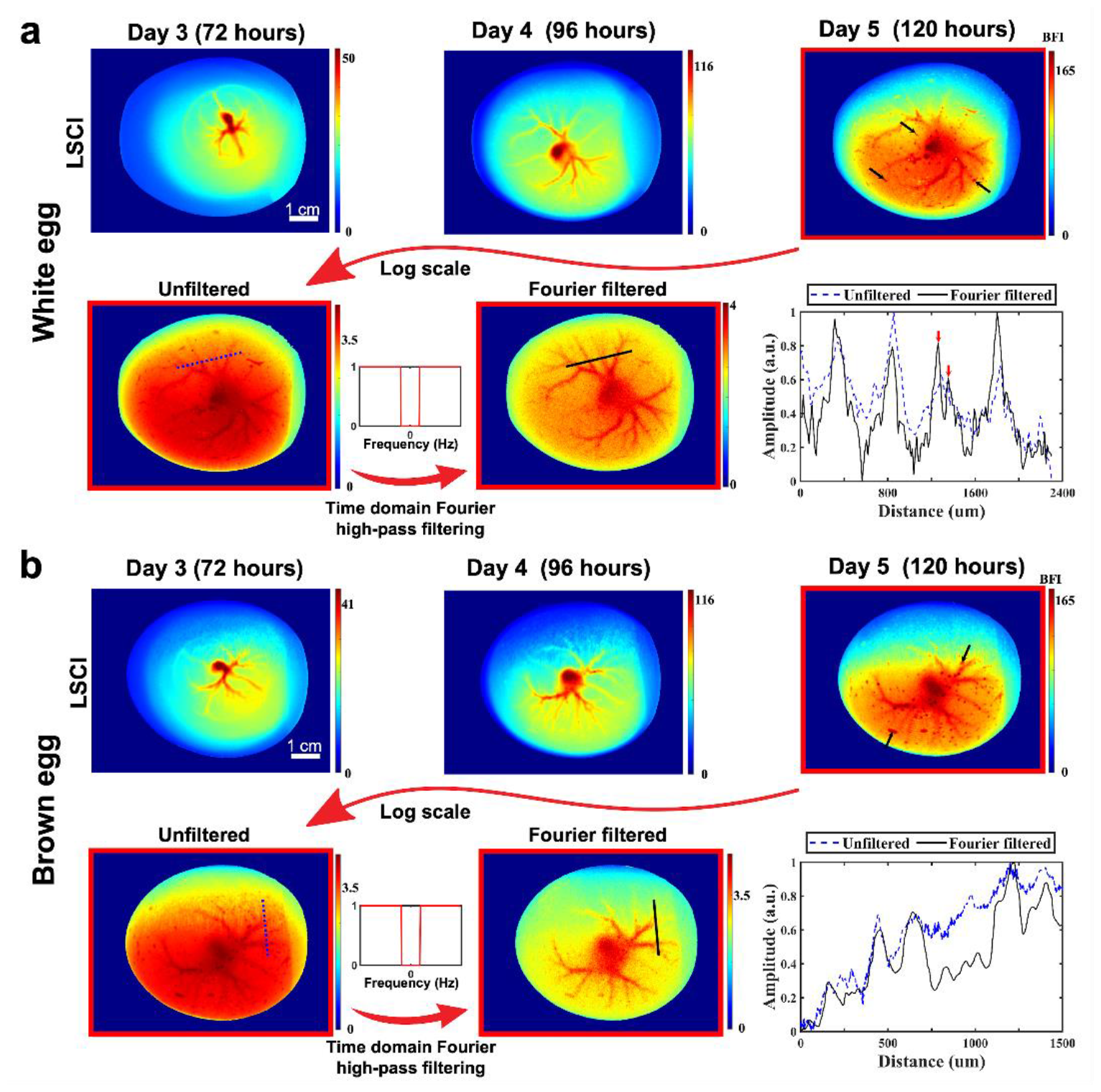
Blood vessel network imaging at different early days of incubation: day 3 (72 hours), day 4 (96 hours), and day 5 (120 hours). **a** White egg imaging results. **b** Brown egg imaging results. The imaging artifacts on day 5 from shell cracks and body movements are removed by using a time domain Fourier high pass filtering method. Selected blood vessels cross-section at day 5 are compared in log scale before and after artifact removal.

We non-invasively monitored the blood flow and measured the heartbeat of chicken embryos on day 5, suggesting a comprehensive assessment of their overall health (Fig. 6, Supplementary Movie). The ADF denoiser with a sliding window in time domain facilitated dynamic imaging and heartbeat estimation, significantly enhancing the temporal resolution. Further enhancement in capturing the cardiac cycle could be achieved with a higher frame rate for the recording camera.

**Fig. 6.**
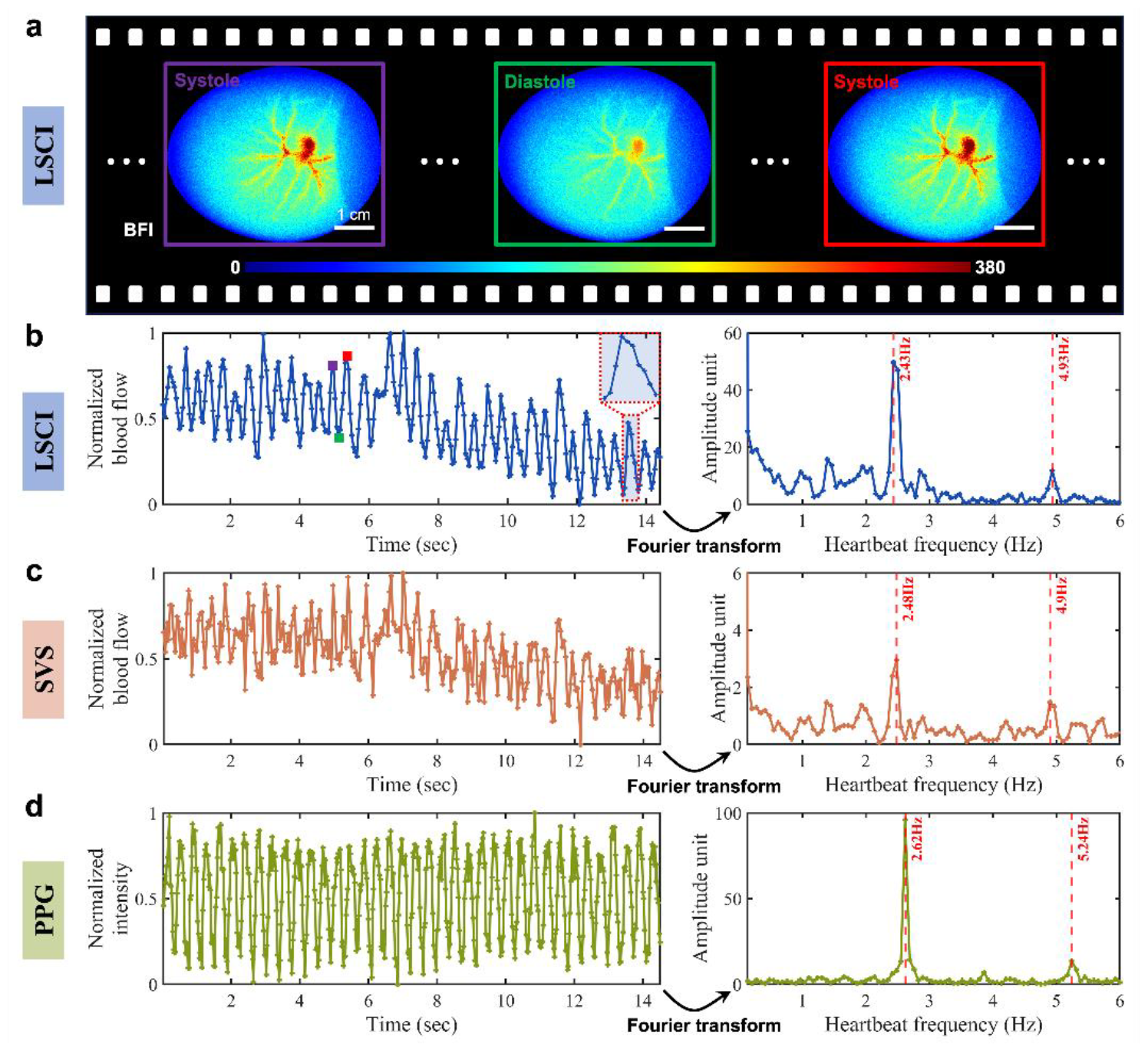
Monitoring the blood flow of a chicken egg at day 4.75 (114 hours) of incubation. **a** Snapshot images from an LSCI movie capturing the systolic and diastolic periods of a cardiac cycle of the chicken egg. **b** Normalized blood flow temporal dynamics calculated from the LSCI method. **c** Normalized blood flow temporal dynamics calculated from the speckle visibility spectroscopy (SVS) method. **d** Normalized intensity calculated from microscope camera images after opening the egg (Photoplethysmography (PPG) method). The Fourier transform plots compare the frequency estimation of heartbeat and its second harmonic among the three methods.

Importantly, we achieved 85% accuracy in classifying the developmental stages of chicken embryos from HH15 to HH22 according to the Hamilton-Hamburger scale, based on a dataset comprised of 260 blood vessel images from LSCI (Fig. 7). This staging technique relies on the blood vessel characteristics of the chicken embryo rather than the body morphology observed after opening the egg. Indeed, even with the egg opened, achieving a staging accuracy of 100% is challenging due to non-uniform body development. For instance, the head may develop faster than the tail, and vice versa. In contrast, blood vessels exhibit a more linear trend of development, making them easier to quantify. We believe that with a larger training dataset, the use of LSCI non-invasive imaging for staging can potentially surpass the accuracy achieved through traditional staging method which involves opening the egg.

**Fig. 7.**
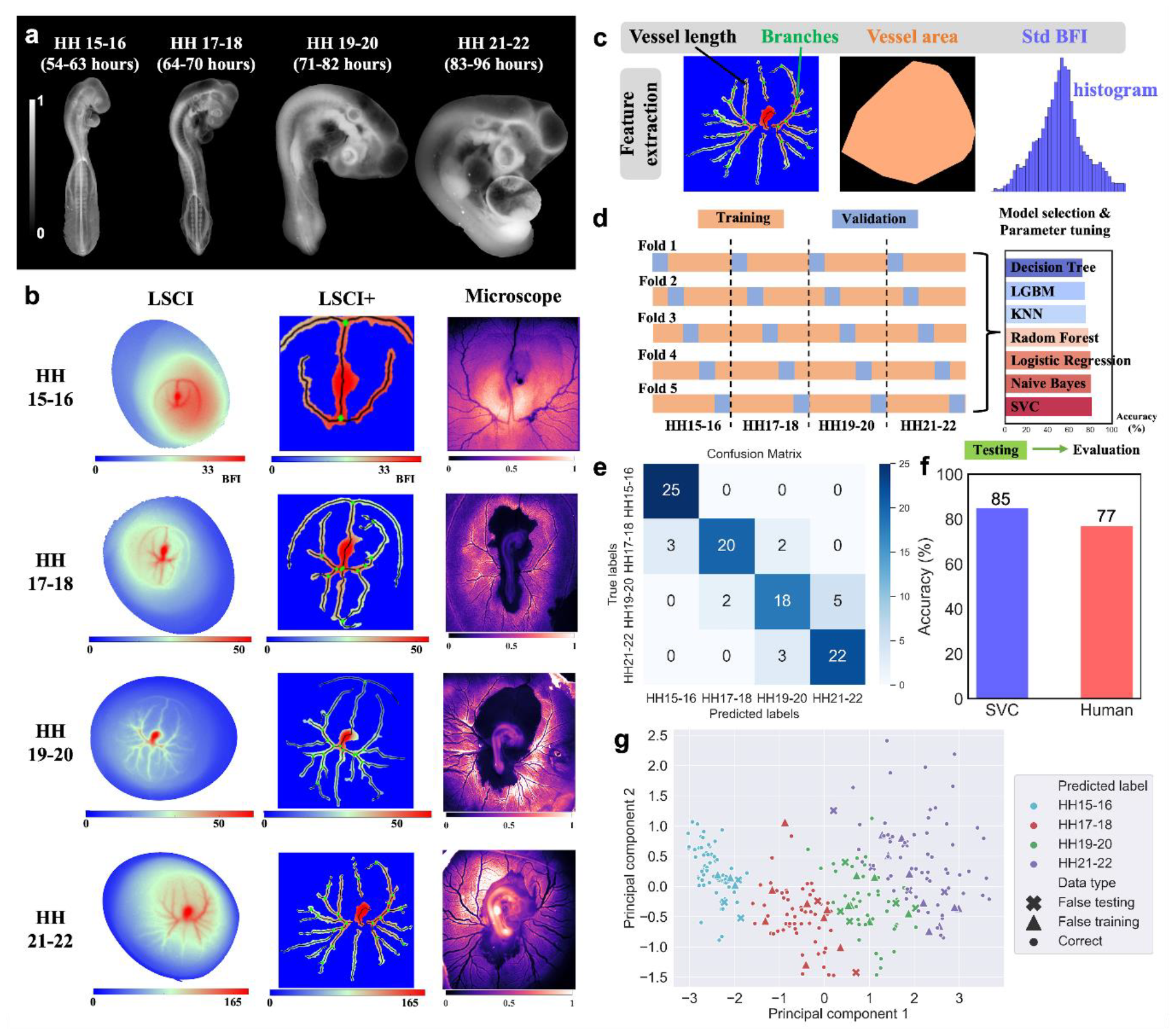
LSCI staging of chicken embryos. **a**. Examples of four Hamburger-Hamilton (HH) stage periods (HH15-16, HH17-18, HH19-20, HH21-22) captured with a wide field microscope. **b**. Typical LSCI images, LSCI+ and microscope images of the CAM at these four stage periods. LSCI+ is an enhanced version of LSCI showing blood vessel skeletons, branches, and BFI value distributions. **c**. Feature extraction process from LSCI+ images. **d**. Dataset splitting for training, validation, and testing. Five-fold cross-validation was used to select the best model and parameters among seven classifiers. Testing and evaluation were done on a separate dataset unseen before. **e**. Confusion matrix of the Support Vector Classifier (SVC). **f**. Classification accuracy comparison between the machine learning algorithm (SVC) and human using LSCI images. **g**. Visualization of feature space distribution and classification performance using SVC. The four features were expressed by the first two principal components using principal component analysis.

In addition to the staging application, we introduced the LSCI+ method (Fig. 7b, c) to non-invasively quantify blood vessel features, including vessel length, area, the number of branches, and blood flow index distributions. As shown, microscope images portray the entire vascular network, including vessels with anomalously low flows, while LSCI selectively highlights large vessels with regular and robust blood flow, showing the potential of the LSCI method for detecting malfunctioning or inactive blood vessels.

From a biological standpoint, our work opens avenues for further exploration, particularly in the quantification of the CAM, heart development study, and the potential for additional non-invasive staging/longitudinal studies. This research not only helps to improve our understanding of avian embryonic development but also holds potential implications for non-invasive sexing of chicken embryo at early-stages of their development.

## Methods

### LSCI image processing

The first step of our processing is to do the noise removal. The intensity value *I* received on the camera can be decomposed as^42,43^:

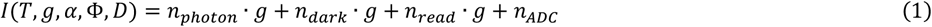

where *T* represents the exposure time, *g* the camera gain, *α* the quantum efficiency, Φ the incident photon flux and *D* the dark current. The photon detection is described by a Poisson distribution of mean *T* . *α* . Φ (*n*_*photon*_∼*Poisson*(*T* . *α* . Φ)), the dark noise as *n*_*dark*_∼*Poisson*(*T* . *D*), the readout noise as *n*_*read*_∼*Normal*(0, *σ*_*read*_), and the analog-to-digital convertor noise as *n*_*ADC*_∼*Normal*(0, *σ*_*ADC*_). The expectation of the mean intensity value over time < *I* >_*t*_ can be written as:

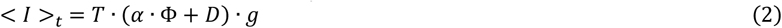

Hence, the dark noise can be estimated by averaging *N* = 200 camera frames captured with no laser exposure (Φ = 0) and the same exposure time *T* with the raw speckles in Fig. 1b. By subtracting dark noise from the raw speckles, we get the dark noise-free intensity 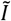:

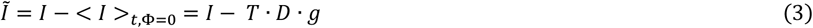

The local speckle contrast value *K* of a speckle image 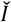 is defined as

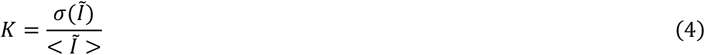

Where 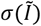 is the standard deviation of the intensity 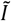, and 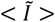 is the mean value of that pixel. Speckle contrast can be calculated either temporally or spatially^17,18^, denoting as tLSCI and sLSCI. sLSCI works on a single speckle frame, in which a sliding window of size 5 × 5 or 7 × 7 is used to calculate local speckle statistics *σ*_*s*_ and *μ*_*s*_. The average <>_*s*_ denotes spatial averaging of all *N*_*s*_ pixels

within the sliding window region.

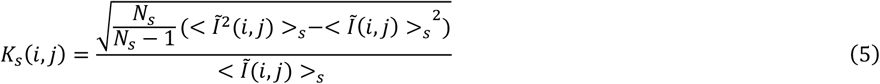

Single sLSCI image is quite noisy, therefore, temporal averaging of *K*_*s*_ is usually used to improve the signal-to-noise ratio, which is called spatial-temporal LSCI (stLSCI).

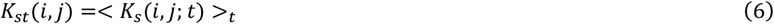

For tLSCI, the temporal speckle contrast *K*_*t*_ can be calculated from a series of *N* speckle pattern images recorded by the camera as^26^:

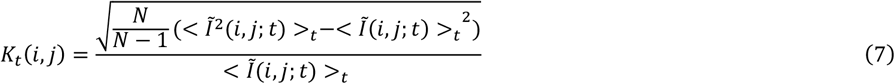

where *σ*_*t*_ is the standard deviation and *μ*_*t*_ is the temporal mean of the *N* recorded speckle pattern images. The intensity at a pixel *i, j* (*i* for row and *j* for column) is denoted 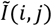, and *t* denotes the time at which the speckle pattern was recorded. The average <>_*t*_ denotes temporal averaging occurring at each pixel *i, j*, see yellow boxes in Fig. 1b. tLSCI is resistant to static scattering because *σ*_*t*_ will not change when adding a static term to 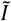. To reduce the dot-like speckle noise after tLSCI reconstruction, we used an anisotropic diffusion filter (ADF)^27,28^, which works in an iterative way by:

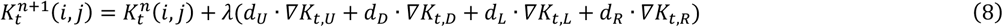

where *n* is the number of filtering iterations, *λ* = 1/4 represents the diffusion rate. *d* and ∇*K*_*t*_ are the diffusion coefficient and the contrast gradient in the direction of four neighboring pixels (up, down, left, right). ADF mimics the heat diffusion process, in which the edges of blood vessels act as restriction boundary conditions. We used the *imdiffusefilt* function in Matlab to implement this processing step and the maximum iteration is set as 30.

Assume that the scattered light is an ergodic field with the absence of static scattering, the Siegert relation holds^48^. Typically, we consider an unordered motion of blood cells and a Lorentzian blood velocity distribution, and the speckle contrast *K* can be related to decorrelation time *τ*_*c*_ of the dynamic movement as^17,36,42^:

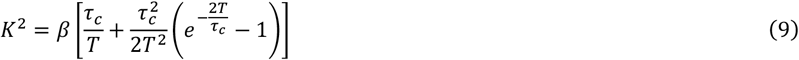

where *β* factor here is a coefficient that accounts for contrast loss due to light polarization, laser coherence, speckle size, and camera sampling^35,37,49^. When *T* ≫ *τ*_*c*_, the second term in equation (9) can be ignored, and it can be simplified as:

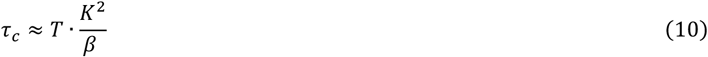

Finally, the blood flow index (BFI) is used to assess blood flow, which can be calculated as the reciprocal of *K* squared^36^.

The faster the blood flow, the shorter the decorrelation time and thus the larger the BFI.

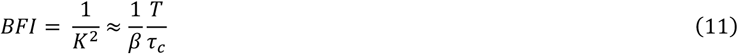

The speckle size of the speckle field using our LSCI setup was calculated by fitting the radial averaged autocorrelation profile within center speckle regions, which results in an estimation of *s* = 1.31 pixels in full width at half maximum. Note that we also proposed the method to remove artifacts in LSCI images, i.e., the Fourier high pass filtering method, more details can be found in Supplementary Fig. 4.

### Dual imaging platform

The schematic of our experimental system is presented in Fig. 1c. The brightfield transmission imaging system is illuminating the egg from its bottom, perpendicularly to the LSCI system. The illumination is a 530 nm LED [Thorlabs M530L4] delivering a 35 nm bandwidth incoherent light with an output power of 480 mW. Such LED can be digitally triggered. A coupling lens [Thorlabs AC254-040-A] is inserted after the LED to collimate the LED light onto the egg. The green light then propagates inside the chicken egg, as shown in Fig. 1a. The transmission light and fluorescence emission exiting the chicken egg is collected by an imaging system and imaged onto the camera.

For the LSCI system, we used a single-longitudinal-mode, single frequency continuous wave 852 nm collimated laser source [Spectra-Physics DL852-300-SO]. This highly coherent laser source is thermally stabilized and can deliver up to 230 mW, useful for laser speckle imaging. The laser light is illuminating the chicken eggs from its side with a beam diameter of 5 mm. The choice of near-infrared (852 nm) wavelength is motivated by the fact that biological tissue penetration is more effective with longer wavelengths^16,17^, but to ensure good quantum efficiency with the camera, we limited our LSCI illumination wavelength to near-infrared. A digitally controlled optical shutter is placed after the laser source and allows for switching on/off the LSCI acquisition. A pair of mirrors in a face-to-face 90 deg configuration is used to precisely direct the light onto the chicken egg.

The chicken egg is inserted on an egg-holder in a horizontal orientation. We designed the egg-holder with an ellipse shape to fit in and stabilize the egg position during imaging. A linear polarizer is used prior to the egg to remove any deleterious polarized light that can cause speckle disruptions with our imaging. The scattered laser light exiting the chicken egg is collected from top view by an imaging system onto the camera.

The imaging system consists first of a mirror changing the propagation orientation from vertical to horizontal. An aperture is inserted next to the mirror to control the region-of-interest of the field-of-view and to filter out undesired stray light. An optical shutter is placed in front of the camera and can be digitally controlled. The optical shutter is used as a filter switch. We used a low-pass filter [Thorlabs FESH0700] in brightfield transmission imaging to only let visible light pass through and a band-pass filter [Thorlabs FBH850-10] of 850±5 nm in the LSCI channel imaging. A camera (including a lens of 50 mm focal length [Edmund Optics #86-574]) records the emitted light from the chicken egg. The size of the aperture near to pupil plane of this imaging lens as well as the magnification of the optical system are optimized together to both ensure field-of-view coverage of the whole egg and the speckle size on the camera plane to be larger than two pixels. In our setup, the magnification is 0.2 and the numerical aperture is 0.03.

For our study, we used a RGB camera [Thorlabs CS126CU] of exposure time ranging from 0.03 ms to 14.7 s. The camera resolution is 3000 × 4096 pixels, with a pixel pitch size of 3.45 × 3.45 µm. The RGB channels of the camera have the same quantum efficiency at 852 nm, ensuring that there is no resolution loss when using it for LSCI imaging. For the brightfield transmission imaging experiments, the camera was in the RGB mode, and with an exposure time set in the range 0.5 s to 3.5 s, depending on the eggshell absorption loss. For the LSCI imaging, the camera was operating in the monochromatic mode with an exposure time set at T = 10 ms and at a speed of 21 frame per second. All the instrument controls were implemented using Matlab and Arduino.

### Dataset collection for staging

The LSCI images were collected according to our designed incubation schedule of 54-63 (HH15-16), 64-70 (HH17-18), 71-82 (HH19-20) and 83-96 (HH21-22) hours. For best imaging, the eggs were rotated to a position where the heart of the vessel distribution is close to the center of the image, using the live view imaging ability of our LSCI system. The eggs were positioned on the sample holder for 2 minutes before imaging to guarantee stabilization of the blood vessel position within the egg. We also kept the top surface of the egg as much as a flat plane as possible to reduce defocus effect. After we finished the LSCI imaging, we opened the egg and captured a microscope image of the chicken embryo. Finally, the microscope images were sent to two experienced biologists for staging ground truth labeling. We used a commercial wide-field microscope (Olympus MVX10 with 1x objective lens) for imaging the embryo body and the vessels (Fig. 7b, Microscope images). To increase the contrast for visualizing the embryo, insoluble black ink was injected under the embryo’s CAM. In each paired stage period, the dataset comprised exclusively with images within the corresponding paired stages. In cases where the actual stage fell between two of the paired stage periods, an experienced biologist will decide the more likely corresponding paired stage period, by principally relying more on morphological differences influencing the blood vessel network development of the embryo, such as the heart and the head.

### Feature extraction from blood flow images

We started from LSCI images and applied some image processing steps (See Supplementary Fig. 5) to get the LSCI+ images (Fig. 7b). Four features were then automatically extracted and quantified as four numbers. The vessel length (in mm) was calculated as the total length of vessel skeleton excluding the outer ring. The number of branches equals to the number of branch points plus number of isolated regions. Vessel area (in mm^2^) includes all the pixel counts within the convex hull of the vessels. Normalized standard deviation of BFI was calculated based on its histogram, which is its standard deviation divided by its mean.

### Incubation of chicken eggs

The animal research for this study received confirmation from the Caltech Office of Laboratory Animal Resources. In all our studies, chicken eggs were incubated no later than day 12. The chicken eggs were incubated using commercial incubators (from Hethya, and Kebonnixs companies) regulating the humidity within the 50-70% range and auto-regulating the temperature within 99.5-100.5°F range. The incubators also had an automatic egg turning function, every 60 or 90 minutes, gently rolling the eggs. The disposal of the chicken egg embryo was performed in compliance with the Caltech Institutional Animal Care and Use Committee policy.

## Data availability

The data that support the findings of this study are available from the corresponding author upon reasonable request.

## Acknowledgements

The authors thanks Caltech FSRI undergraduate students Angelica Moussambote, Said M. Garcia, and Yingyin Tan for their help and assistance during the imaging experiments. The authors also thanks professor Jerome Mertz, graduate students Yu Xi Huang, Siyu (Steven) Lin, Haowen Zhou and Ruizhi Cao for fruitful discussions and advice. This research was supported by the IST Carver Mead Endowment — Award No. 25550038. Simon Mahler is the recipient of the 2024 SPIE-Franz Hillenkamp Postdoctoral Fellowship.

## References

1. Gandhi, S., Ezin, M. & Bronner, M. E. Reprogramming Axial Level Identity to Rescue Neural-Crest-Related Congenital Heart Defects. Developmental Cell 53, 300-315.e4 (2020).

2. Goenezen, S., Chivukula, V. K., Midgett, M., Phan, L. & Rugonyi, S. 4D subject-specific inverse modeling of the chick embryonic heart outflow tract hemodynamics. Biomech Model Mechanobiol 15, 723–743 (2016).

3. Burggren, W. & Rojas Antich, M. Angiogenesis in the Avian Embryo Chorioallantoic Membrane: A Perspective on Research Trends and a Case Study on Toxicant Vascular Effects. JCDD 7, 56 (2020).

4. Nowak-Sliwinska, P., Segura, T. & Iruela-Arispe, M. L. The chicken chorioallantoic membrane model in biology, medicine and bioengineering. Angiogenesis 17, 779–804 (2014).

5. Waddington, C. H. III. Experiments on the development of chick and duck embryos, cultivated in vitro. Phil. Trans. R. Soc. Lond. B 221, 179–230 (1932).

6. New, D. A. T. A New Technique for the Cultivation of the Chick Embryo in vitro. Development 3, 326–331 (1955).

7. Bronner-Fraser, M. Methods in Avian Embryology. (Academic Pr, San Diego, Calif., 1996).

8. Sukumaran, V. et al. Experimental assessment of cardiovascular physiology in the chick embryo. Developmental Dynamics 252, 1247–1268 (2023).

9. Pi, S. et al. Angiographic and structural imaging using high axial resolution fiber-based visible-light OCT. Biomed. Opt. Express 8, 4595 (2017).

10. Galli, R. et al. Sexing of chicken eggs by fluorescence and Raman spectroscopy through the shell membrane. PLoS ONE 13, e0192554 (2018).

11. Informatics in Poultry Production: A Technical Guidebook for Egg and Poultry Education, Research and Industry. (Springer Nature Singapore, Singapore, 2022).

12. Jia, N. et al. A Review of Key Techniques for in Ovo Sexing of Chicken Eggs. Agriculture 13, 677 (2023).

13. Hall, C. A., Potvin, D. A. & Conroy, G. C. A new candling procedure for thick and opaque eggs and its application to avian conservation management. Zoo Biology 42, 296–307 (2023).

14. Ernst, R. A., Bradley, F. A., Abbott, U. K., Craig, R. M. & Craig, R. M. Egg Candling and Breakout Analysis. (University of California, Agriculture and Natural Resources, 2004).

15. Jia, N. et al. Exploratory Study of Sex Identification for Chicken Embryos Based on Blood Vessel Images and Deep Learning. Agriculture 13, 1480 (2023).

16. Draijer, M., Hondebrink, E., Van Leeuwen, T. & Steenbergen, W. Review of laser speckle contrast techniques for visualizing tissue perfusion. Lasers Med Sci 24, 639–651 (2009).

17. Boas, D. A. & Dunn, A. K. Laser speckle contrast imaging in biomedical optics. J. Biomed. Opt. 15, 011109 (2010).

18. Dunn, A. K., Bolay, H., Moskowitz, M. A. & Boas, D. A. Dynamic Imaging of Cerebral Blood Flow Using Laser Speckle. J Cereb Blood Flow Metab 21, 195–201 (2001).

19. Bandyopadhyay, R., Gittings, A. S., Suh, S. S., Dixon, P. K. & Durian, D. J. Speckle-visibility spectroscopy: A tool to study time-varying dynamics. Review of Scientific Instruments 76, 093110 (2005).

20. Senarathna, J., Rege, A., Li, N. & Thakor, N. V. Laser Speckle Contrast Imaging: Theory, Instrumentation and Applications. IEEE Rev. Biomed. Eng. 6, 99–110 (2013).

21. Dunn, A. K. Laser Speckle Contrast Imaging of Cerebral Blood Flow. Ann Biomed Eng 40, 367–377 (2012).

22. Li, P., Ni, S., Zhang, L., Zeng, S. & Luo, Q. Imaging cerebral blood flow through the intact rat skull with temporal laser speckle imaging. Opt. Lett. 31, 1824 (2006).

23. Du, E., Shen, S., Chong, S. P. & Chen, N. Multifunctional laser speckle imaging. Biomed. Opt. Express 11, 2007 (2020).

24. Padmanaban, P. et al. Assessment of flow within developing chicken vasculature and biofabricated vascularized tissues using multimodal imaging techniques. Sci Rep 11, 18251 (2021).

25. Chen, R., Miao, P. & Tong, S. Transmissive multifocal laser speckle contrast imaging through thick tissue. J. Innov. Opt. Health Sci. 16, 2350005 (2023).

26. Yang, L. et al. Noninvasive vasculature detection using laser speckle imaging in avian embryos through intact egg in early incubation stage. Biomed. Opt. Express 4, 32 (2013).

27. Sang, X., Chen, B., Li, D., Pan, D. & Sang, X. Transient Thermal Response of Blood Vessels during Laser Irradiation Monitored by Laser Speckle Contrast Imaging. Photonics 9, 520 (2022).

28. Song, L. et al. Improving temporal resolution and speed sensitivity of laser speckle contrast analysis imaging based on noise reduction with an anisotropic diffusion filter. J. Opt. 20, 075301 (2018).

29. Hamburger, V. & Hamilton, H. L. A series of normal stages in the development of the chick embryo. Journal of Morphology 88, 49–92 (1951).

30. Preuße, G. et al. Highly sensitive and quick in ovo sexing of domestic chicken eggs by two-wavelength fluorescence spectroscopy. Anal Bioanal Chem 415, 603–613 (2023).

31. Gillbro, T. & Cogdell, R. J. Carotenoid fluorescence. Chemical Physics Letters 158, 312–316 (1989).

32. Goodman, J. W. Speckle Phenomena in Optics: Theory and Applications. (Roberts, Englewood, Colo, 2007).

33. Chriki, R. et al. Spatiotemporal supermodes: Rapid reduction of spatial coherence in highly multimode lasers. Phys. Rev. A 98, 023812 (2018).

34. Mahler, S. et al. Fast laser speckle suppression with an intracavity diffuser. Nanophotonics 10, 129–136 (2020).

35. Briers, J. D. Laser speckle contrast analysis (LASCA): a nonscanning, full-field technique for monitoring capillary blood flow. J. Biomed. Opt. 1, 174 (1996).

36. Liu, C., Kiliç, K., Erdener, S. E., Boas, D. A. & Postnov, D. D. Choosing a model for laser speckle contrast imaging. Biomed. Opt. Express 12, 3571 (2021).

37. Yordanov, S., Drucker, M.Butt, H.-J. & Koynov, K. Real-time monitoring of biomechanical activity in aphids by laser speckle contrast imaging. Opt. Express 29, 28461 (2021).

38. Kirkpatrick, S. J., Duncan, D. D. & Wells-Gray, E. M. Detrimental effects of speckle-pixel size matching in laser speckle contrast imaging. Opt. Lett. 33, 2886 (2008).

39. Duncan, D. D., Kirkpatrick, S. J. & Wang, R. K. Statistics of local speckle contrast. J. Opt. Soc. Am. A 25, 9 (2008).

40. Phuphanin, A., Sampanporn, L. & Sutapun, B. Smartphone-Based Device for Non-Invasive Heart-Rate Measurement of Chicken Embryos. Sensors 19, 4843 (2019).

41. Mahler, S. et al. Assessing depth sensitivity in laser interferometry speckle visibility spectroscopy (iSVS) through source-to-detector distance variation and cerebral blood flow monitoring in humans and rabbits. Biomed. Opt. Express 14, 4964 (2023).

42. Xu, J., Jahromi, A. K. & Yang, C. Diffusing wave spectroscopy: A unified treatment on temporal sampling and speckle ensemble methods. APL Photonics 6, 016105 (2021).

43. Huang, Y. X., Mahler, S., Mertz, J. & Yang, C. Interferometric speckle visibility spectroscopy (iSVS) for measuring decorrelation time and dynamics of moving samples with enhanced signal-to-noise ratio and relaxed reference requirements. Opt. Express 31, 31253 (2023).

44. Hertzman, A. B. THE BLOOD SUPPLY OF VARIOUS SKIN AREAS AS ESTIMATED BY THE PHOTOELECTRIC PLETHYSMOGRAPH. American Journal of Physiology-Legacy Content 124, 328–340 (1938).

45. Groves, I. et al. Accurate staging of chick embryonic tissues via deep learning of salient features. Development 150, dev202068 (2023).

46. Stern, C. D. Staging tables for avian embryos: a little history. Int. J. Dev. Biol. 62, 43–48 (2018).

47. Martinsen, B. J. Reference guide to the stages of chick heart embryology. Developmental Dynamics 233, 1217–1237 (2005).

48. Carpenter, D. K. Dynamic Light Scattering with Applications to Chemistry, Biology, and Physics (Berne, Bruce J.; Pecora, Robert). J. Chem. Educ. 54, A430 (1977).

49. Wang, Y. et al. Improving the estimation of flow speed for laser speckle imaging with single exposure time. Opt. Lett. 42, 57 (2017).

